# Interactions between xylem traits linked to hydraulics during xylem development optimize growth performance in conifer seedlings

**DOI:** 10.1101/2021.12.24.474017

**Authors:** Jehová Lourenço, Daniel Houle, Louis Duchesne, Daniel Kneeshaw

## Abstract

- Climate change has threatened forests globally, challenging tree species’ ability to track the rapidly changing environment (e.g., drought and temperature rise). Conifer species face strong environmental filters due to climatic seasonality. Investigating how conifers change their hydraulic architecture during xylem development across the growing season may shed light on possible mechanisms underlying hydraulic adaptation in conifers.
- Laser microscopy was used to assess the three-dimensional hydraulic architecture of balsam fir (*Abies balsamea* (Linnaeus) Miller), jack pine (*Pinus banksiana* Lambert), white spruce (*Picea glauca* (Moench) Voss), and black spruce (*Picea mariana* (Miller) Britton, Sterns & Poggenburgh) seedlings. We measured hydraulic-related xylem traits in different regions (from early to latewood), during four years of plant growth.
- The xylem development of jack pine seedlings contrasts with the other species by maintaining torus overlap (a hydraulic safety-associated xylem trait), relatively constant across the season (from early to latewood), and over the years studied. The expansion of tracheids and torus was positively associated with plant growth.
- Pit aperture-torus covariance is central to the seasonal dynamics of jack pine xylem development, which jointly with a rapid tracheid and pit expansion is consistent with strong growth performance. Linking xylem structural changes during xylem development with hydraulics is a major issue for future research to assess conifers vulnerability to climate change.

## Introduction

Climate change has resulted in concomitant increases in temperature and drought (IPCC, 2021), which are particularly strong at high latitudes (e.g. boreal forest regions) (Gauthier *et al*., 2015). Considering that drought is a major environmental driver of tree mortality worldwide (Allen *et al*., 2010), the survival of trees in a climate-change context may rely upon their capacity to modify their hydraulic architecture during xylem development (xylogenesis) to track the rapidly changing environment (Anderegg *et al*.,2015; Guérin *et al*., 2020). Sensitivity to drought has been associated with a tradeoff in plants between hydraulic safety (small conduits, that increase the ability to resist embolism formation and spread) or efficiency (large conduits, increasing the water transport capacity and vulnerability to drought) (Dixon & Joly, 1895; van der Sande *et al*., 2019). However, many plant species from regions with important climatic seasonality seem to depart from the proposed hydraulic tradeoff (Liu *et al*., 2020), suggesting that modifications in the xylem hydraulic traits during xylogenesis should confer the ability to adapt and grow in such changing environments. Thus, the investigation of how multiple hydraulic-related xylem traits change throughout xylogenesis (i.e., from early to latewood) of plants adapted to climatic seasonality may shed light on the mechanisms by which conifers adapt the hydraulic architecture in response to the environmental change to optimize growth along with the growing season.

Xylem tissue it’s an important component to assess plant-environment relationships, as it provides the anatomical basis for the plant hydraulic functioning (Sperry *et al*.,2002) and structural stability (Niklas *et al*., 2006), reflecting different resource-use and growth strategies (Chave *et al*., 2009). In this context, wood anatomy measures the maximum theoretical hydraulic conductivity and carbon allocation priorities at the moment of xylem formation (Guérin *et al*., 2020), including multiple xylem traits (e.g., interconduit pits, parenchyma, and fibers) potentially interacting with conduits to contribute to the water transport (Bittencourt *et al*., 2016; Jupa *et al*., 2016; Venturas *et al*., 2017; Klein *et al*., 2018). For example, an augmented water transport capacity via large conduits and pits has been associated with high photosynthetic capacity and growth performance in plants (Poorter *et al*., 2010; Roskilly *et al*., 2019).

However, the high complexity of the xylem also provides multiple facets to support hydraulic safety. Modifications to pit density (number of pits per unit of tracheid wall volume) are likely a way to achieve this result (Jasińska *et al*., 2015). Pit density has been negatively correlated to xylem-specific hydraulic conductivity, and it is hypothesized to reflect the same variation in conduits (e.g., tracheid density, given by the number of tracheids per xylem area), where the increase in the number of small conduits and pits should provide additional resistance to drought (Jasińska *et al*., 2015). Thus, pit density, such as conduit density (Lens *et al*., 2011; Von Arx *et al*., 2012), is likely a complementary strategy to increase hydraulic integration via interconduit connectivity. Similarly, the number of tracheids connected laterally by pits (here referred to as pit-tracheid net size, *NS*) should increase the hydraulic connectivity. Despite xylem’s integration has been linked to hydraulic pathway redundancy and the xylem’s ability to manage the spread of embolism (Lens *et al*., 2011; Von Arx *et al*., 2012), there is no simple consensus on the significance of xylem integration for drought stress resilience, as it may simultaneously increase the probability of embolism spread between conduits during drought stress and facilitate hydraulic failure. Considering its potential hydraulic role, xylem integration deserves further investigation. Analysis of the coordination between multiple xylem traits (e.g., tracheids and pits) along with the xylem development and its linking to the species growth performance across the season may provide some clues and guidance for future research on this matter.

In this context, intertracheid pits are critical components of the hydraulic system of conifer species, given their important role in xylem hydraulic integration (Venturas *et al*., 2017). The bordered pit is a chamber that contains a permeable membrane, whose central portion has a thickened and impermeable torus. The pit functions as a valve that must deal with the conflict of allowing water flow between functional tracheids while preventing airflow from embolized tracheids by sealing the pit aperture (*Da*) with the torus (Hacke *et al*., 2004) when embolism occurs (Delzon *et al*., 2010). Bordered pits can account for over 50% of the total hydraulic resistance in the xylem, playing an important function in the balance between safety and efficiency in the vascular transport of conifers (Choat *et al*., 2008). Greater dimensions of pit aperture (*Da*) and pit membrane (*Dm*) are hypothesized to compensate for high hydraulic resistance in unicellular and short tracheids, arguably making the hydraulic system of conifers as efficient as the multicellular and large vessels of angiosperms (Pittermann *et al*., 2005).

Thus, any anatomical change in pit traits properties and dimensions during xylogenesis is expected to have profound implications for a plant’s hydraulic functioning, likely playing important adaptative roles in conifers. For example, the permeability of the pit membrane (margo porosity) reduces the flow resistance of xylem sap through pits, increasing the xylem hydraulic efficiency (Schulte *et al*., 2015). The pit membrane and torus flexibility are important features for pit-level hydraulic safety, as they are related to the torus sealing the pit aperture without the rupture of the pit membrane (Schulte *et al*.,2015; Schulte & Hacke, 2021) and the torus (Zelinka *et al*., 2015). The scaling relationships between pit traits dimensions, in turn, should permit fine hydraulic adjustments (Hacke *et al*., 2004) that could be central for plants to adapt and grow under steep environmental changes driven by seasonality (Liu *et al*., 2020). Proportional changes in pit traits should hypothetically guarantee high hydraulic conductivity through pits (e.g., large *Da*) and embolism resistance (large torus overlap, *O*) (Hacke *et al*., 2004), whereas a loose control of this mechanism, should cause plants to tradeoff between hydraulic efficiency (e.g. high *Da/Dt* ratio) and safety (e.g. low *Da/Dt* ratio) over the growing season.

The wide variety of functional traits potentially influencing plant fitness (e.g., growth performance) emphasizes the point that plant responses to seasonality-driven environmental change during xylogenesis may not be achieved by modifications in a single functional trait (Venturas *et al*., 2017). Instead, phenotypic plasticity and adaptation during xylogenesis are expected to be the outcome of a larger number of traits interacting in a more complex network. Investigations into these interactions may shed light on the structural mechanisms underlying such important biological processes as plant hydraulic functioning and growth strategies. High plasticity in hydraulic-related xylem traits may allow boreal tree species to optimize growth by taking advantage of higher soil water due to snowmelt (from early to mid-growing season) in warmer periods (Zhang *et al*., 2019), and also respond to declines in soil water as the growing season progress (Houle *et al*., 2012). High plasticity would also permit boreal seedlings to rapidly change the hydraulic architecture to unexpected droughts due to climatic anomalies (Gauthier *et al*., 2015).

Here, we measured a suite of hydraulic-related xylem traits from early to latewood of jack pine, balsam fir, black and white spruce seedlings, in four different growth rings (2016-2019). We used network analysis to investigate the central traits driving the seasonal dynamics in xylem development, which may shed light on how conifer seedlings change their hydraulic architecture in response to the strong environmental filters driven by climatic seasonality. We specifically addressed the following hypothesis: 1) The xylem traits variability from early to latewood of the studied species will reflect in possible mechanisms linked to hydraulic safety and efficiency, allowing plants to cope with the hydrological changes driven by climatic seasonality; 2) Seedling with both hydraulic efficiency (larger *TLA* and *Dm*) and safety-related xylem traits (e.g., larger *Dt* and *O*) are expected to have better growth performance; 3) tree seedlings should vary between having a tight vs. loose control on pit trait variability resulting in ecological strategies that contrast between stabilizing vs. trading off the variability in pit traits linked to hydraulic safety (e.g., *O*); and 4) we expect that pit traits will be central to the network of interactions in the seasonal dynamics of hydraulic-related xylem traits.

## Materials and Methods

### Study site

Seedlings of balsam fir (*Abies balsamea* (Linnaeus) Miller), jack pine (*Pinus banksiana* Lambert), white spruce (*Picea glauca* (Moench) Voss), and black spruce (*Picea mariana* (Miller) Britton, Sterns & Poggenburgh) were pre-cultivated in a tree nursery and subsequently transplanted to the study area in June of 2014, following a standard distance of 40 cm from each other. This experimental plantation is located in the balsam fir and white birch bioclimatic domain at the Montmorency forest, approximately 50 km north of Québec City, Canada (47°25N, 71°19W, 800-840 m above the sea). The site is a cut block of a mature fir stand harvested with the protection of regeneration and soil in the summer of 2013, following a hemlock looper (Lambdina fiscellaria (Guenée)) outbreak. The site is characterized by a slope of about 12% that is oriented to the north. The soil is a mesic podzolic sandy till of ortho-humo-ferric type. The average temperature (1980-2010) is −0.3 °C and the average annual precipitation is 1665 mm.

### Wood anatomy analysis

In the fall of 2019, we sampled wood in the main seedling stem (10cm above the soil surface) from 10 individuals per species. Wood samples of approximately 4 cm^2^ were transversally sliced with a microtome at 40 microns thickness. The wood microsections were, then, placed in slides, and one drop of immersion oil was added over it. Afterward, we placed coverslips over the microsections and sealed the coverslips boarders with nail polish to produce semipermanent slides. Then, the microsections were scanned using a confocal microscope (Nikon A1 plus, lasers *Alexa 488 water* and *Cy5*) with an objective of 60x of magnification, aperture of 1.40, and diffractive index of 1.51 (Plan apo λ 60x oil). On average, we scanned 100 stacks of 0.175 μm thickness, each with one-third overlap among stacks as this provides a more accurate three-dimensional reconstruction of the wood anatomy architecture. We scanned four different tree rings (2016 to 2019) and three different regions of the wood: 1) <u>earlywood</u>: region of the first-formed cells; 2) <u>intermediate</u>: transition region cells in the median position of the tree-ring; and 3) <u>latewood</u>: region of the last formed cells (See Fig. S1 – supplementary material). This was done to evaluate whether seasonal changes were consistent or variable between years. In total, we produced 388 images, including jack pine (n=103), balsam fir (n=71), white spruce (n=97), and black spruce seedlings (n=117). The numbers of images produced per species diverge because the xylem cells of some wood samples did not have good integrity.

### Xylem traits

We assessed the wood hydraulic architecture by measuring several hydraulic-related xylem traits. The measurements were made in three different regions (earlywood, intermediate, and latewood). We made five measurements of pit diameter, torus size, pit aperture size, tracheid wall thickness, and length, per tree-ring region. These measurements were posteriorly used to calculate the torus overlap (*O*, proportion of torus covering pit aperture), the margo space (*Msp*, pit chamber space not covered by the torus) (Hacke *et al*., 2004; Roskilly *et al*., 2019), and the tracheid-wall reinforcement given by the ratio between the thickness of the double tracheid wall and its span [(t/b)^2^] (Hacke, 2001). Moreover, we measured the tracheid lumen area (*TLA*), tracheid density (*TD*), pit density (*PD*, number of pits per volume), and tracheid-pit net size (*NS*, number of tracheids connected laterally by pits). All tracheids (116 ± 37) of the scanned area (45,220 μm^2^) were accounted for these measurements. For the NS measurements, we accounted for all tracheids-pit connections found in the volume of wood analyzed (630,000 μm^3^). The measurements were performed in the software ImageJ, version 1.52p (Wayne Rasband National Institutes of Health, USA). A detailed description of the trait calculations and hydraulic functions can be found in table 1.

**Table 1.** Overview of wood anatomy traits, their acronym, units, a short description regarding the calculation methods and function, and reference underlying the trait concept.

| Acronym | Unity | Trait | Description | Function | Reference |
| --- | --- | --- | --- | --- | --- |
| TLA | $\mu\text{m}^2$ | Mean tracheid lumen area | Size of the conductive area of the tracheid through which water is transported. | Hydraulic conductivity. | (Carlquist, 2001) |
| TWR | unitless | tracheid wall reinforcement | $(t/b)^2$, where “t” is the double cell wall thickness and “b” the length of the same cell wall. | Hydraulic safety and structural support. Increases tracheid wall hydraulic resistance against implosion and reinforces xylem structure. | (Hacke *et al.*, 2001) |
| TD | $\text{no.mm}^{-2}$ | Tracheid density | Number of tracheids per squared millimeter. | Increases hydraulic integration via pathway redundancy. | (Espino & Schenk, 2009; Cai & Tyree, 2010) |
| Dt | $\mu\text{m}$ | Torus diameter | Pit torus diameter. | Embolism resistance. Protects pit valves against failure. | (Hacke *et al.*, 2004) |
| Da | $\mu\text{m}$ | Pit aperture diameter | Pit aperture diameter. Resistance to the water flow. | Intertracheid hydraulic efficiency. | (Hacke *et al.*, 2004) |
| Dm | $\mu\text{m}$ | Pit diameter | Pit chamber diameter. Resistance to the water flow. | Intertracheid hydraulic efficiency. | (Roskilly *et al.*, 2019) |
| PD | $\text{no.mm}^{-3}$ | Pit density | The number of pits per cubic millimeter. | Intertracheid hydraulic efficiency. | (Jacobsen *et al.*, 2016) |
| O | Unitless | Torus overlap | $O = (Dt - Da)/(Dm - Da)$. Hacke *et al.*’s formula is relativized to pit membrane minus pit aperture diameter, making the calculation more relevant to water flow resistance. | Embolism resistance. Covering pit aperture. | (Hacke *et al.*, 2004; Roskilly *et al.*, 2019) |
| Msp | Unitless | Margo space | $M = (Dm - Dt)/(Dm - Da)$. It accounts for the distance between the pit membrane border (m) and torus (t) relativized to the pit membrane minus pit aperture diameter (a). | Hydraulic efficiency. It represents the space through which water must flow in the pit. | (Hacke *et al.*, 2004; Roskilly *et al.*, 2019) |
| NS | Unitless | Pit-tracheid net size | Mean size of the tracheid-pi-/tracheid net. Larger nets have a greater number of tracheids connected laterally. We analyzed a wood volume of $630,000 \mu\text{m}^3$ per 3D scanned image. | Hydraulic efficiency and safety. Increases hydraulic conductivity. | |

### Linear discriminant and correlations analysis

Before the linear discriminant analysis (LDA), the data was standardized/normalized, using estimate processing parameters. This is a recommended procedure, as discriminant analysis can be affected by the predictors’ measurement units (James *et al*., 2014). The models (seasonal plasticity, early and latewood) were fitted by the *lda* function of the MASS R package (Venables & Ripley, 2002), reaching accuracies above 0.72 (Table 2). The term “seasonal plasticity” is used here to define the variance (or changes) in xylem traits that occurred within the growing season, and was assessed by the calculation of the variance on the xylem traits mean values measured from early to latewood, in each studied tree-rings/years (2016 to 2019). This procedure reduced the data points to 133 observations (annual mean and variance of trait values), including jack pine (n=36), balsam fir (n=28), white spruce (n=29), and black spruce (n=40). The ordination plots were made with the *ggord* R package version 1.1.1. (Beck, 2020). The species were also compared regarding the correlation and scaling relationship between the pit traits and between pit traits and tracheid lumen area (*TLA*). We used the *cor.test* R function to calculate the correlations, while *lm* R function was used for the linear models that were built to extract the slope (or scaling relationship) between the predictor and the response variable (e.g., *O* ~ Da). The R scripts of the LDA calculation can be found in the online version of this article.

**Table 2.**
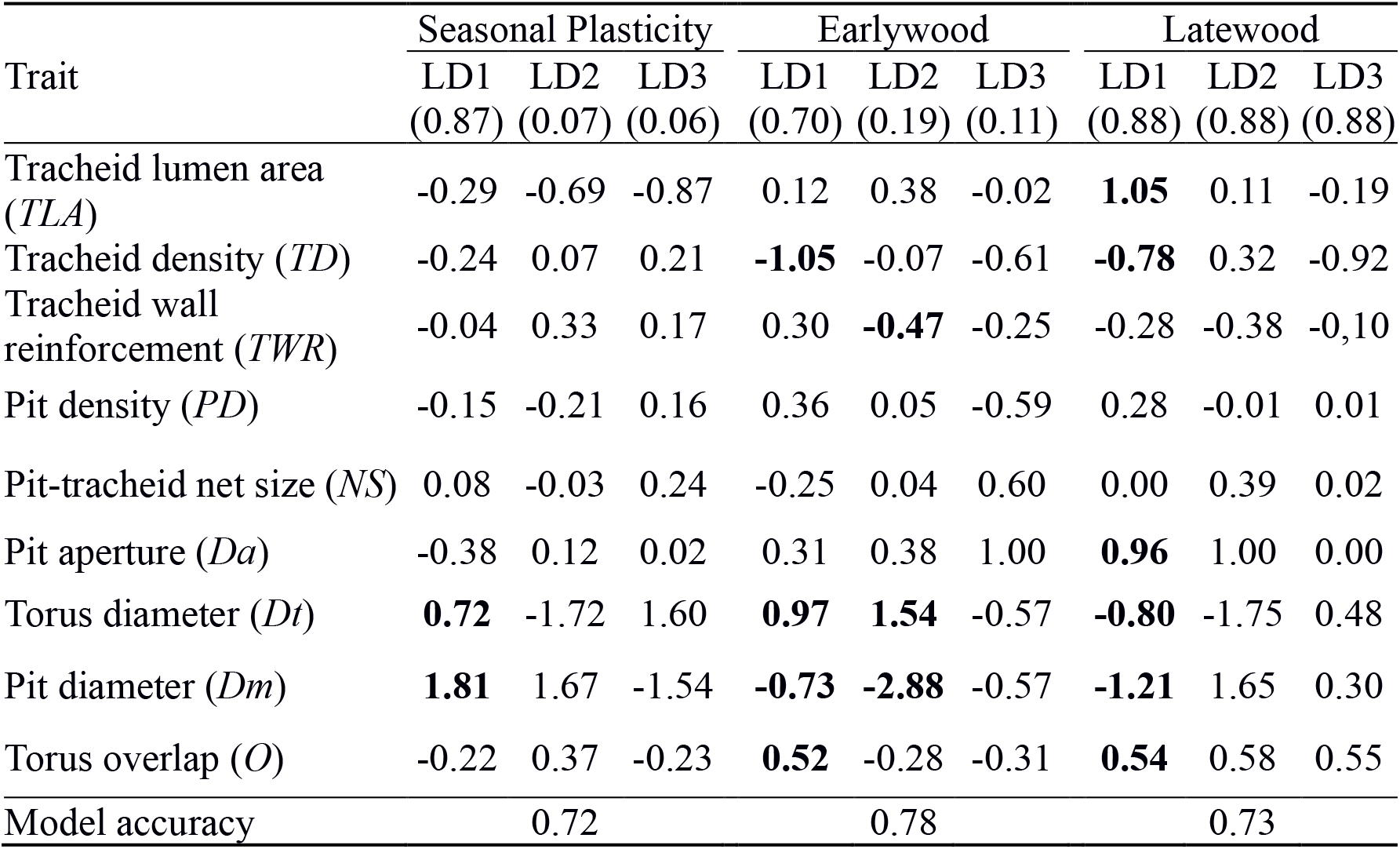
Coefficients of linear discriminants based on jack pine (JKP), black spruce (BSP), white spruce (WSP), and balsam fir (BFR) hydraulic-related xylem traits. The proportion of explained variation for each ordination axis of the linear discriminants (LD) is shown within the brackets. Seasonal plasticity was calculated by the variance of the trait values measured from early to latewood, i.e. across the growing season. The highest values are highlighted in bold.

### Network analysis

Network analysis is a promising tool to untangle biological complexity (Poorter *et al*., 2014) such as in plant hydraulic functioning. Its usage has been extended to the investigation of trait integration and functional differentiation among co-existing plant species (Burton *et al*., 2020) as well as to unravel the role of photosynthetic and hydraulic tradeoffs (Poorter *et al*., 2014; De La Riva *et al*., 2016), providing a way to effectively represent plant functioning and phenotypic plasticity (Messier *et al*., 2017).

Network analysis is based on the premise that the interaction between the components in a system is key to understanding the whole system’s functioning (Csardi & Nepusz, 2006), and it is used here to test the importance of pit traits to the observed seasonal plasticity in the xylem/hydraulic architecture of conifer seedlings. It is a quantitative approach to graphically represent and investigate complex systems given the interaction of their components, which is based on their connectivity and distances. The interactive components, hereafter xylem traits, are represented by nodes and their connectivity, i.e., correlations, are represented by the edges linking them. The size of the nodes and edges represent the strength of trait centrality regarding the whole network and the strength of the correlation between a pair of traits, respectively. The R function *as.undirected* was used to create undirected edges. Network graphics were constructed and the betweenness centrality analysis was made in the *igraph* R package (Csardi & Nepusz, 2006), using the *betweenness* function to test which traits are central to hydraulic adjustments of the studied plant species in response to environmental changes within the season.

The betweenness value gives the number of shortest paths from all nodes to all others passing through the focal node, which reveals the importance of the node for the network, here graphically represented by the node size. A similar conceptual trait network (Fig. 4a) was adopted for all species, however, only correlations of 0.5 and above were used in the analysis, to highlight the more relevant trait correlations (Fig. S2, for details). We used the algorithm Kamada-Kawai for the graphic layout, which has been largely used in network analysis of plant functioning (Poorter *et al*., 2014; De La Riva *et al*.,2016; Messier *et al*., 2017; Burton *et al*., 2020). The algorithm finds the bidimensional optimal position of the traits in a network, reflecting their relative correlation strengths and thus the traits interactions that may influence the hydraulic safety and efficiency of the xylem across the season.

**Figure 4.**
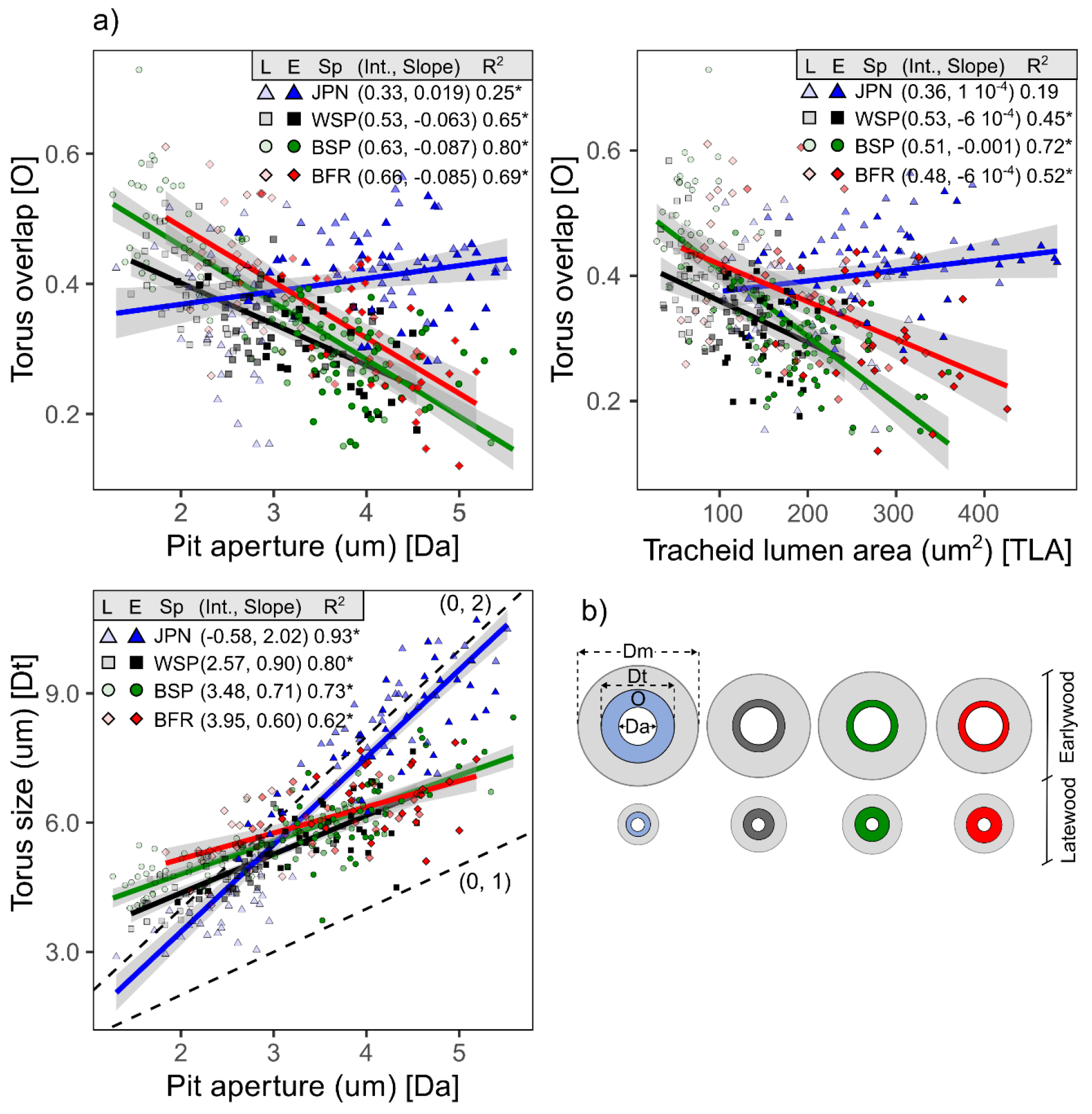
(a) Relationships between the variation in pit aperture and tracheid lumen area with the variation in torus overlap in seedlings of white spruce (WSP), black spruce (BSP), jack pine (JPN), and balsam fir (BFR). The intercept (Int.), slope, and Pearson correlation of the regression lines are shown. (b) The scheme represents pit trait variability in species resulting from their different scaling relationships. The asterisks highlight statistically significant correlations (p-value<0.05). Lighter dots represent the xylem latewood (L) and darker dots xylem earlywood (E). Da – pit aperture; Dt – torus size; and O – torus overlap.

## Results

### Species differences in annual and seasonal variation in hydraulic-associated xylem traits

The comparison of species traits shows that jack pine stands out for its high plasticity (or high seasonal variance in the mean trait value) in torus size (*Dt*) and pit diameter (*Dm*) (longer arrows/vectors in Fig. 1a). However, all species substantially shift in xylem trait space, from early to latewood (Fig. 1b,c). In the earlywood, hydraulic-related xylem traits are similar between species, as they markedly overlap in trait space (Fig. 1b). In the latewood, jack pine diverges from the other species by having a larger tracheid lumen area (*TLA*, Fig. 1c, Fig. 2). Seedlings of jack pine had the largest measured TLA across years and growth ring regions (Fig. 2). This species also maintained larger pit aperture (*Da*), *Dt*, *Dm*, and *torus overlap* (*O*) until intermediate regions of the xylem development over the years, while *Dm* was relatively larger in the latewood of the other species (Fig. 2). Jack pine seedlings also had the best growth performance achieving up to 2 meters height at the end of 2019. In general, the growth of all studied species is positively associated with the enlargement of *TLA*, *Dm*, and *Dt* (Fig. 3).

**Figure 1.**
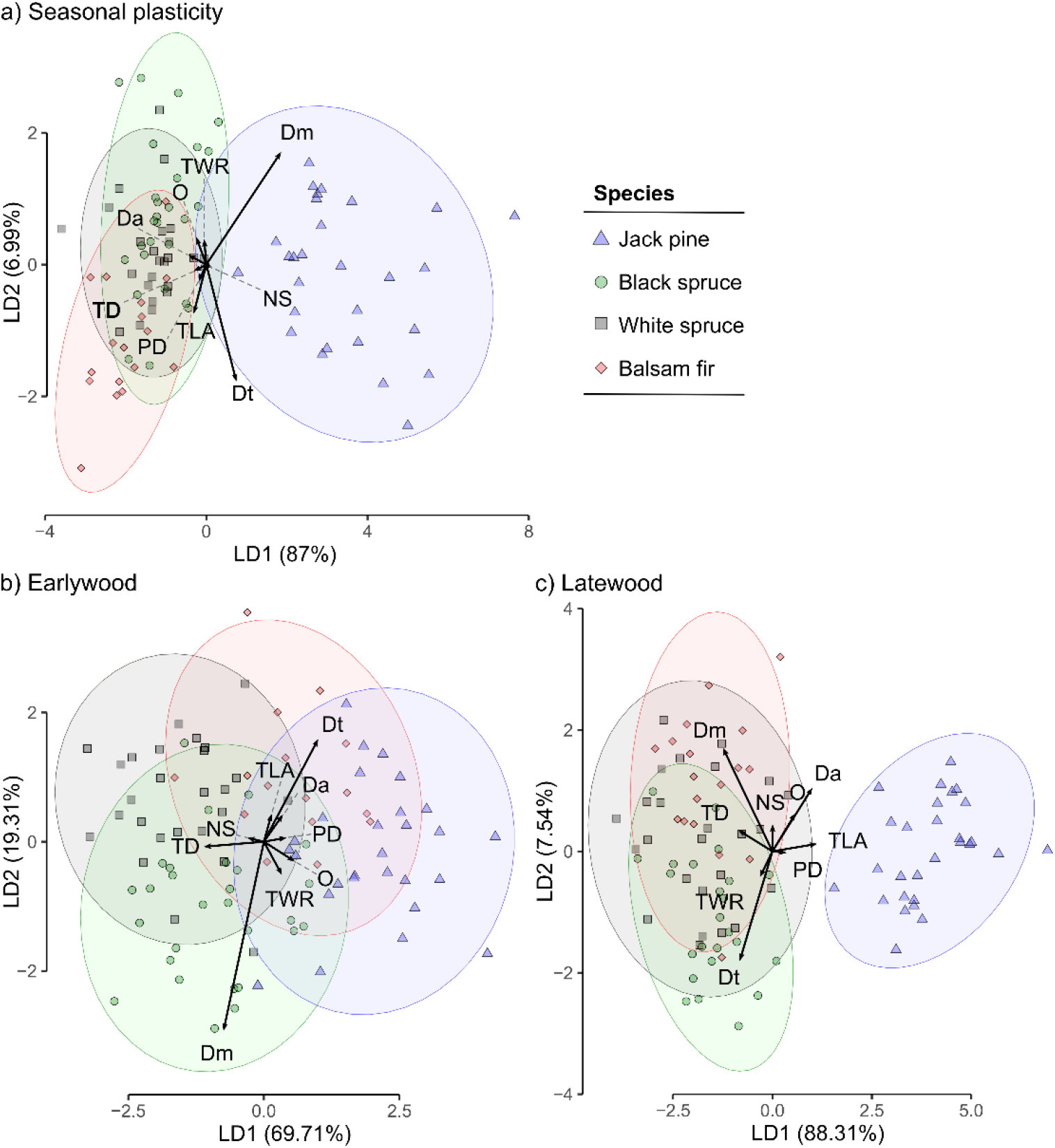
Linear discriminant analysis showing species differences in (a) xylem traits plasticity (mean trait value variance from early to latewood) across the growing season, mean values of xylem traits in the (b) earlywood, and (c) latewood. Jack pine remarkably differs by having the highest seasonal plasticity in torus (*Dt*) and pit diameter (*Msp*), whereas white spruce, black spruce, and balsam fir overlap in the ordination space, reflecting lower seasonal plasticity in pit traits. Traits are torus overlap (*O*), tracheid lumen area (*TLA*), tracheid wall reinforcement (*TWR*), tracheid density (*TD*), torus diameter (*Dt*), pit diameter (*Dm*), pit density (*PD*), pit aperture (*Da*), mean pit-tracheid net size (NS). Dashed lines were included to link small vectors to their names. A detailed description of traits can be found in Table 1.

**Figure 2.**
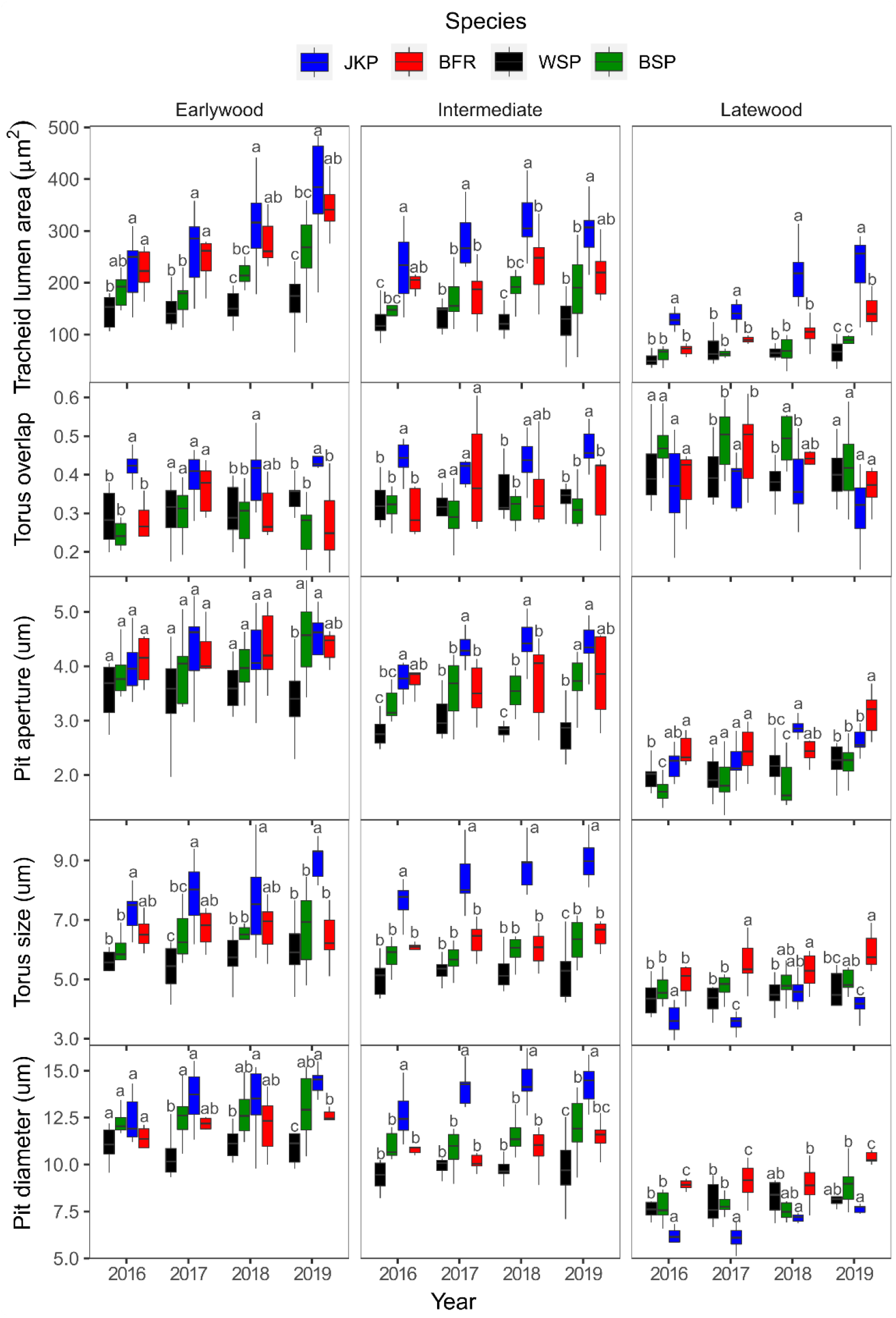
(a) Boxplots showing variability in xylem traits of white spruce (WSP), black spruce (BSP), jack pine (JPN), and balsam fir (BFR), within different regions in the wood (earlywood, intermediate, and latewood) from 2016 to 2019. Letters indicate statistical differences (p-value<0.05) between species compared within the same year and growth-ring region. The test was computed via Tukey’s “honest significant difference” method.

**Figure 3.**
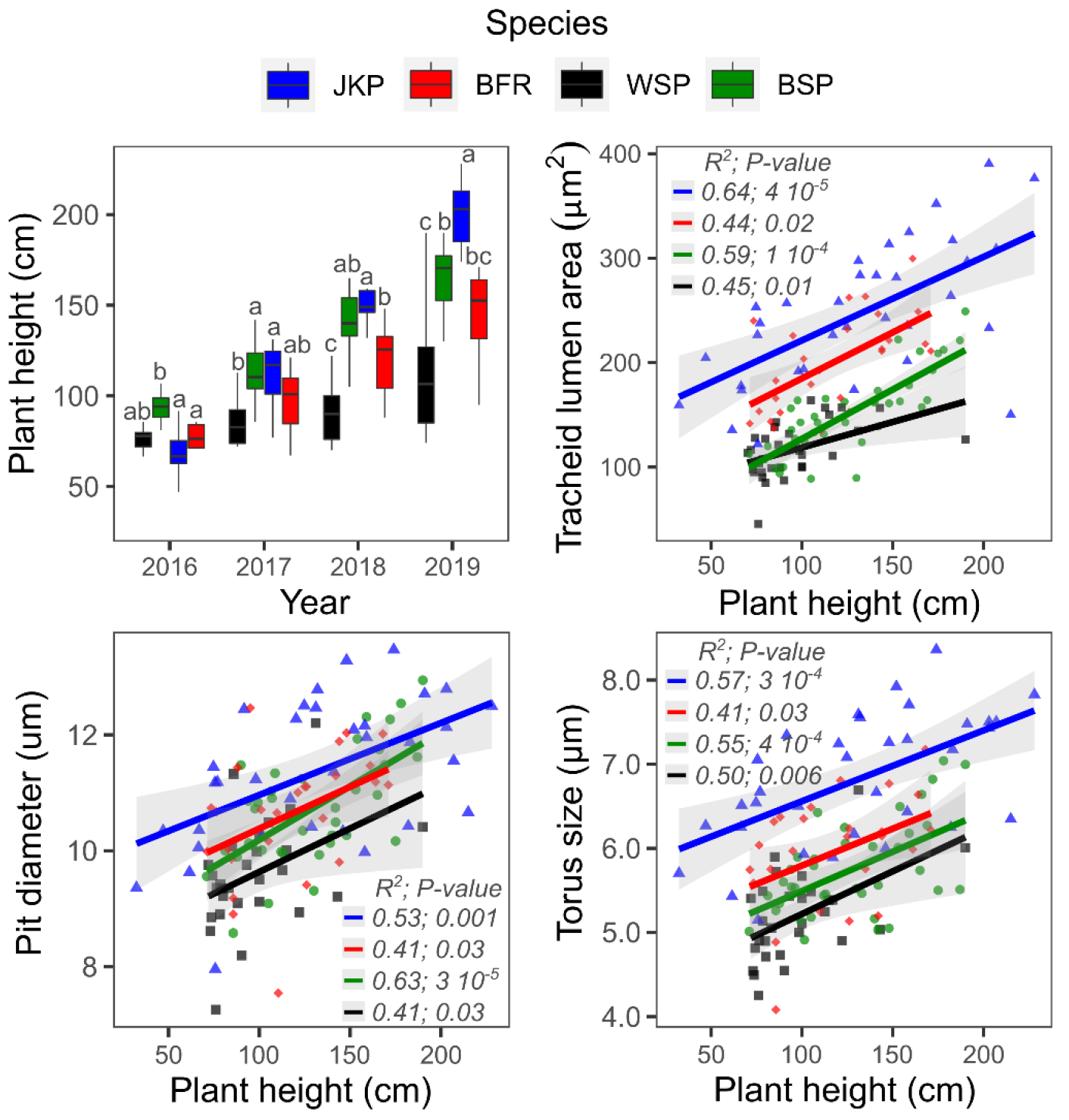
Boxplot showing variation in plant height across the years (2016-2019), and scatter plots showing the relationships between increasing plant height and xylem traits of white spruce (WSP), black spruce (BSP), jack pine (JPN), and balsam fir (BFR). In the scatter plots, each point is an annual mean trait value calculated from the measurements made across different regions in the wood (from early to latewood).

### Scaling relationship between xylem traits and variability in torus overlap (O)

The high seasonal plasticity (variance in the mean trait value within the season) in pit diameter (*Dm*) and torus size (*Dt*) (Fig. 1a) allowed jack pine to maintain high and stable values of torus overlap (*O*, ~0.4) across the growing season (Fig. 3a). This resulted in the largest pits in the earlywood (Fig. 3b). Although highly correlated to pit aperture (*Da*), the torus size of jack pine changed twice as fast as *Da* (Fig. 3a, r^2^ = 0.93 and slope = 2.02), allowing this species to maintain the relative proportions of pit traits that affect *O* (Fig. 2b). In contrast, the other species varied widely in *O* (0.30 to 0.60), due to the lower correlation and regression slope between *Dt* and *Da* (e.g., balsam fir, r^2^=0.62 and slope=0.60). Hence, balsam fir, black and white spruce seedlings tradeoff between low vs. high *O* in the early vs. latewood (Fig. 3b). Except for jack pine, the increase in *TLA* is followed by a decrease in *O* (Fig. 3a). Lastly, the scaling relationship between *Dt* and *Da* was stable across years in the xylem of all species (Table S2 – see the supporting information for additional results).

### Trait network interactions and tradeoffs

The network analysis shows that species diverge in network centrality (Table 3 and Fig. 5b) which can be attributed to seasonal variability (i.e., changes from early to latewood) in torus size (*Dt*) and pit aperture (*Da*) (jack pine); pit diameter (*Dm*) (black spruce); *Dm* and pit-tracheid net size (*NS*) (balsam fir); *Da* and *Dm* (white spruce). Pit and tracheid enlargement have conflicting relationships with *O* in balsam fir, black and white spruce, which is not observed for jack pine *O*, which is only consistently associated with *Dt* (Fig. 5b). Thus, except for jack pine, the increase in multiple traits related to hydraulic efficiency (e.g., *Dm*, *Da*, tracheid lumen area (*TLA*), Fig. 5c) leads to conflicting or negative relationships with pit-level hydraulic safety (i.e., *O*). Despite the species network differences, some similar tradeoffs (or negative correlations) were found in tracheid traits (tracheid lumen vs. wall reinforcement vs. density) and between tracheid and pit traits (e.g. tracheid wall reinforcement (*TWR*) vs. *Da* and *Dm*). Also, no tradeoff was found between pit diameter and density, although both are positively correlated to pit-tracheid net size (Fig. 5b). Opposite to tracheid density (*TD*), pit density and *NS* are small in the latewood (Fig. S3). The *O* is more strongly correlated to torus size in jack pine, while it is strongly associated with *Da* in the other species (see Fig. S2 for the complete species matrix of trait correlations).

**Figure 5.**
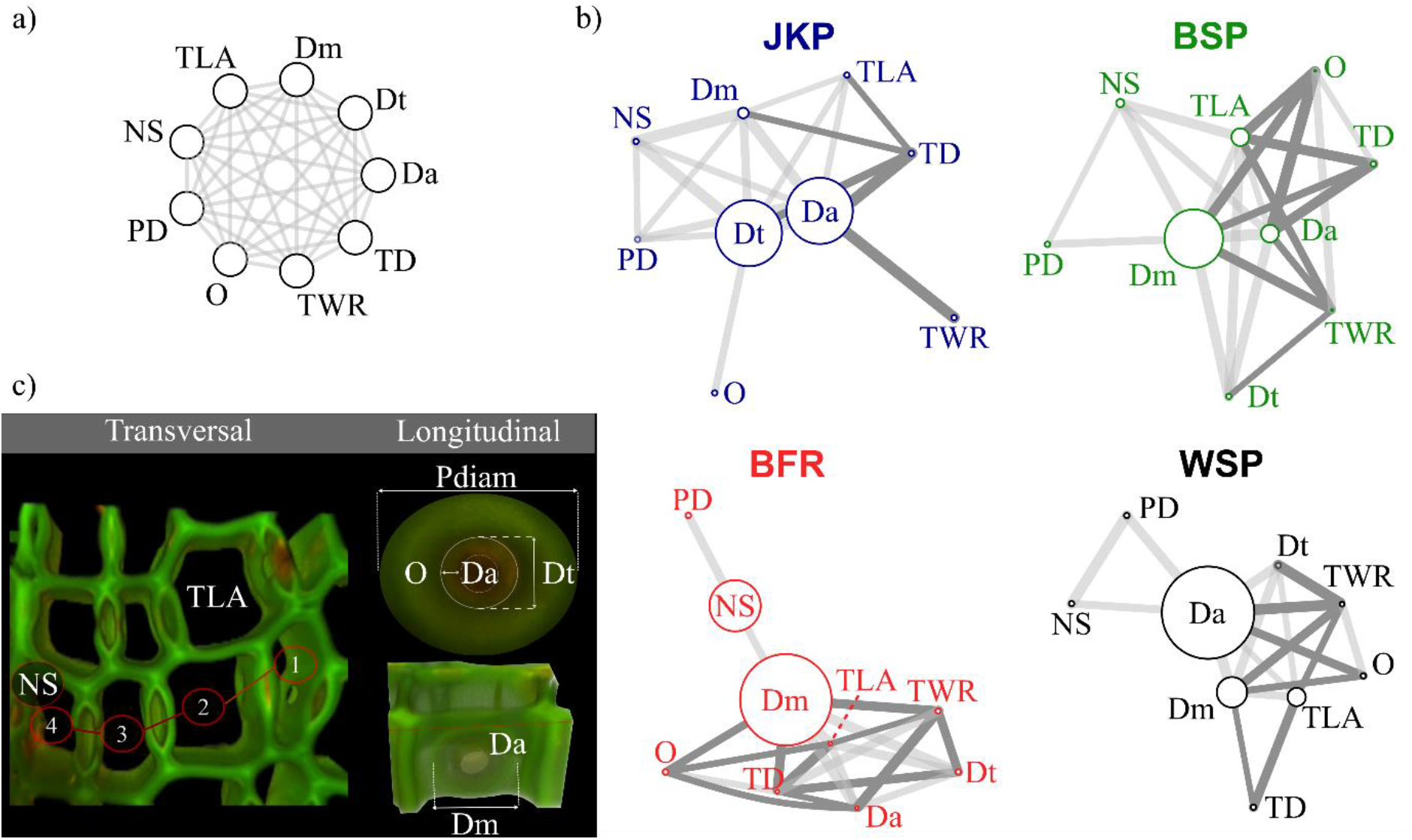
(a) The conceptual network, highlighting all expected correlations between traits, (b) trait correlation networks of jack pine (JKP), white spruce (WSP), black spruce (BSP), and balsam fir (BFR), and (c) three-dimensional images of the xylem traits. The location of traits was optimized by network analysis (igraph, r package) based on the strength of Pearson correlation between traits represented by edge thickness (gray bars) and distance among traits (edges). Only pairs of traits with r^2^≥0.5 and p-value>0.05 were considered (see Fig. S1). Edge colors highlight positive (light gray) and negative (dark gray) correlations among a pair of traits. The vertices sizes are scaled to betweenness centrality, where larger vertices indicate greater betweenness centrality, here interpreted as the importance of a trait to the network of traits interaction and variation along with the growing season. Traits are torus overlap (*O*), tracheid lumen area (*TLA*), tracheid wall reinforcement (*TWR*), tracheid density (*TD*), torus diameter (Dt), pit diameter (*Dm*), pit density (*PD*), pit aperture (*Da*), mean pit-tracheid net size (*NS*). A detailed description of traits can be found in table 1.

**Table 3.** Trait network centrality, considering the seasonal variability in xylem traits of jack pine (JKP), balsam fir (BFR), white spruce (WSP), and black spruce (BSP). The centrality statistic reported is “betweenness”, which considers the path length, i.e., the more edges and the shorter they are, the higher the betweenness value, which is interpreted as the importance of a xylem trait to the changes observed in the xylem. The highest betweenness values are highlighted in bold.

| Trait | Centrality |  |  |  |
| --- | --- | --- | --- | --- |
|  | JKP | BSP | WSP | BFR |
| Tracheid lumen area (*TLA*) | 0 | 2.16 | 2.50 | 0.50 |
| Tracheid density (*TD*) | 0 | 0 | 0 | 0.50 |
| Tracheid wall reinforcement (*TWR*) | 0 | 0.25 | 0.66 | 0 |
| Pit density (*PD*) | 0 | 0 | 0 | 0 |
| Pit-tracheid net size (*NS*) | 0 | 1.00 | 0 | 7.00 |
| Pit aperture (*Da*) | **8.33** | 2.16 | **12.66** | 0.50 |
| Torus diameter (*Dt*) | **8.33** | 0 | 0 | 0 |
| Pit diameter (*Msp*) | 1.33 | **7.16** | 4.16 | **12.50** |
| Torus overlap (*O*) | 0 | 0.25 | 0 | 0 |

## Discussion

### Pit traits have the largest seasonal variability in species xylem, while TLA and Dt enlargement is strongly associated with the outstanding growth performance of jack pine seedlings

The high seasonal variation in xylem traits seems to underlie important hydraulic mechanisms, as hypothesized, which may be crucial for plants to adapt to seasonal environmental changes. In particular, the high seasonal variability in the torus diameter (*Dt*) and pit diameter (*Dm*) of jack pine suggests that pit traits should be important components of the wood development of this species, likely conferring better adaptability to the changes in soil water availability, which is highest after snowmelt and decreases as the summer advances. All pit traits of jack pine reduce in size from earlywood to latewood while tracheid lumens (*TLA*) are significantly larger than balsam fir and spruce seedlings throughout the growing season. These findings suggest a strong control of water flow characteristics via pits (Schulte *et al*., 2015) in jack pine. We argue that the quick variation in *Dt* should increase the xylem embolism resistance via torus overlap (*O*) (Hacke *et al*., 2004) in jack pine seedlings, allowing this species to have larger tracheids and pits for a high water transport capacity until later stages of wood development (intermediate regions of the growth ring, Fig. 2).

The differences between species concerning variation in hydraulic efficiency and safety-related xylem traits (especially, TLA and Dt) may explain their differences in growth performances (Fig. 3). Our results suggest that the fast growth of jack pine seedlings is associated with the rapid expansion of tracheids, pits, and torus. The enlargement of conduits and pits is expected to reduce the water flow resistance as the plants grow taller (Olson *et al*., 2020), while *Dt* and *O* should improve the safety of the hydraulic system functioning (Hacke & Jansen, 2009). These traits should result in an increased capacity of transporting water to the leaves for photosynthesis and carbon acquisition (Galmés *et al*., 2007), ultimately favoring plant growth (Roskilly *et al*., 2019).

### Seedlings vary between having a tight vs. loose control on pit aperture and torus size scaling relationships, resulting in a constant vs. variable torus overlap during the xylem development

Our results corroborate the hypothesis that variability in tight vs. loose control of pit traits results in species either maintaining or varying torus overlap (*O*) during the wood development. In this context, jack pine seedlings maintained high and stable values of *O* by tightly coordinated adjustments in pit aperture diameter (*Da*) and torus diameter (*Dt*). These modifications should guarantee high resistance to embolism at the pit level (Hacke *et al*., 2004). On the other hand, weak correlations and scaling relationships between pit traits in balsam fir seedlings could lead to moments of lower resistance to water flow and weaker embolism control by pits (large *Da* and low *O*) in the earlywood, and moments of higher water resistance and greater embolism control by pits (large *Dt* and *O*) in the latewood. These observations suggest the existence of seasonal variation in pit traits dimensions and scaling relationships, particularly between *Dt* and *Da*. However, the reduction of *O* with an increase in *TLA* observed in balsam fir and spruce seedlings could lead to physiological consequences, making the xylem of these species to tradeoff between hydraulic efficiency (large *TLA* and small *O*) and safety (small *TLA* and large *O*) (Pittermann *et al*., 2010).

We argue that a tight scaling between pit aperture and torus size is important for maintaining consistent safety margins of *O*, protecting pit valve functioning against drought-induced torus displacement (Venturas *et al*., 2017). A high *O* should also allow jack pine seedlings to, while large *TLA* and *Da* should optimize hydraulic conductivity (Schulte *et al*., 2015) and plant growth (Roskilly *et al*., 2019). In other words, jack pine’s xylem architecture seems to get around the safety-efficiency tradeoff hypothesis (Dixon & Joly, 1895; van der Sande *et al*., 2019), as large tracheids, pits, and *O* should provide both an efficient and safe hydraulic system. Ultimately, the co-optimization of both safety and efficiency could result in the maintenance of a high proportion of functional tracheids even in older sapwood regions, which could have subsidiary or additive effects on water transport (Utsumi *et al*., 2003; Bryukhanova & Fonti, 2013). The scaling variability in pit traits presented here provides new pathways of research on plant hydraulics and reinforces the need for future investigations to expand the role of pit traits interactions on conifers’ xylem vulnerability.

### Pit traits have a central role in conifers xylem traits network and tradeoffs

Our results corroborate the hypothesis that pit traits are central in the conifer species trait networks and seasonal dynamics during the xylem development. However, species diverge regarding the most central pit trait, suggesting contrasting hydraulic mechanisms or evolutionary solutions in conifers (Bryukhanova & Fonti, 2013) to cope with dynamic environmental changes promoted by seasonal variation (i.e., high moisture after snowmelt followed by generally declining soil moisture throughout the growing season). Our results, showing the centrality of the covariation between pit aperture and torus size in jack pine, are consistent with an interpretation of the central role of pit traits on multiple xylem traits. Such pit trait covariation could provide a way to resolve the conflict between increasing xylem traits related to hydraulic efficiency (e.g. via tracheid and pit enlargement) and embolism resistance (e.g. torus overlap, *O*) (Gleason *et al*.,2015). We argue that this *Da-Dt* coordination could be a possible mechanism underlying the safety-efficiency hydraulic co-optimization predicted for conifers species (Liu *et al*.,2020). Indeed, jack pine trees typically occur in sandy soils with low water retention (Desponts & Payette, 1992; Carcaillet *et al*., 2020; Pacé *et al*., 2020), and a constant embolism resistance (e.g., via *O*) may be beneficial in such environments where soil water is strongly variable.

In contrast, seasonal xylem dynamics of the other species show conflicting relationships (negative correlations) in several hydraulic efficiency-related xylem traits with *O*. The coordination between pit diameter (*Dm*) and tracheid-pit net size (*NS*) is central for balsam fir hydraulic adjustments, and differently than hypothesized (Jasińska *et al*., 2015), increasing pit diameter/net-size/density seems to occur in the earlywood (see Fig. S3), where most of the studied species should be more vulnerable to embolism (low *O*) and be more hydraulic efficient (larger tracheids). Thus, pit-tracheid connectivity and xylem integration seem mostly associated with the increase in hydraulic efficiency in all species, instead of safety. Xylem integration in conifers could increase vulnerability to embolism and hydraulic failure if species have reduced *O*, as seen in the earlywood of balsam fir and spruces seedlings. These results contradict the previous hypothesis (von Arx *et al*., 2013; Jasińska *et al*., 2015), and highlight the need for investigations on pit-tracheid relationships to elucidate the effects of xylem integration on hydraulic efficiency and safety of conifers. Lastly, tracheid wall reinforcement (*TWR*) tradeoffs with *TLA* and pits (*Dm* and *Da*), suggesting that the thickening or the reinforcement of the cells should be primarily associated with xylem hydraulic resistance due to constraining the opening size of pits (Hacke *et al*., 2004) and *TLA* (Schuldt *et al*., 2016). These modifications should reduce the water flow, the pace of carbon assimilation, and plant growth (Chave *et al*., 2009) by the end of the growing season.

Despite our findings suggesting possible linkages between xylem traits and plant hydraulics, we acknowledge that our interpretations are limited to the potential effect of xylem traits on species’ hydraulic safety and efficiency. However, our research is framed under the proposition that wood anatomy guides research on plant hydraulics (Olson *et al*., 2020; Lourenço Jr. *et al*., 2022), providing new perspectives and promising areas of investigation on possible intrinsic factors (anatomical) underlying plants’ ability to cope with environmental changes (e.g., drought and temperature rise). The pit traits and tracheid interaction presented here may ultimately reflect species fitness differences (Violle *et al*., 2007), and thus further investigation may help to predict what species are more likely to survive to the effects of climate change.

## Conclusion

The capacity of gymnosperm pits to transport water while controlling the spread of embolism (Hacke *et al*., 2004; Pittermann *et al*., 2005) and possible interactions with tracheids could provide a wide range of adjustments and configurations in xylem architecture, by which conifer species could increase embolism resistance (e.g., via high torus overlap, *O*) and still have a highly efficient hydraulic system with large tracheids and pits (pit structure and aperture). Our results suggest that conifer species seem to have evolved a tightly coordinated mechanism that adjusts the xylem architecture via pit aperture/torus covariation. Coordination between tracheid and pit traits enlargement could underlie the hypothesized hydraulic safety-efficiency co-optimization in conifers (Liu *et al*., 2020) and plausibly be the outcome of the selective forces acting on the hydraulic systems of conifer species via climatic seasonality. Therefore, we suggest that the investigation of the xylem seasonal dynamics during wood development should be a major issue to understanding conifer adaptability and vulnerability to environmental change. The investigation of the possible physiological implications of the pit-tracheids scaling relationships is a key area of research for advancing our understanding of the intrinsic (anatomical) factors controlling the hydraulic functioning and the growth performance of conifers.

## Acknowledgments

This work was conducted at the University of Quebec in Montreal and JLJ was financed by Mitacs (FR38551). We are grateful for Mitacs and Ouranos financial support. We would like to thank Dr. Loïc D’Orangeville who designed and supervised the installation of the experimental plantation. We also thank the forestry agencies for the cooperation and the field technicians of the Ministry of Forests, Wildlife, and Parks and the numerous students (particularly Éric Brazeau-Goupil) who participated in the implementation and maintenance of the plantation.

## Author Contribution

JLJ, DK, DH, and LD designed the project; JLJ, DH, and LD carried out the sampling; JLJ supervised the preparation and analysis of the samples, analyzed the data; JLJ, DK, DH, and LD led the writing of the manuscript. All authors contributed critically to the drafts and gave final approval for the publication.

## Supporting information

**Figure S1.** Scheme showing the scanned regions within the tree-ring.

**Figure S2.** Species correlation matrices.

**Table S1.** Regression parameters of xylem traits by year.

**Coding.** R scripts for LDA and network analysis.

## Notes

### Competing Interest Statement

The authors have declared no competing interest.

