## Supplementary material for "Interactions between xylem traits linked to hydraulics during xylem development optimize growth performance in conifer seedlings"

Authors: Jehova Lourenco Junior, Daniel Houle, Louis Duchesne, Daniel Kneeshaw.

The following Supporting Information is available for this article:

**Fig. S1** Scheme showing the tree-ring scanned regions.

**Fig. S2** Species correlation matrices.

**Fig. S3** Boxplots showing additional analysis of seasonal and annual variation in pit traits.

**Table S1** Regression parameters of xylem traits by year.

**Fig. S1** Scheme of 3d scanned regions within the tree-rings.

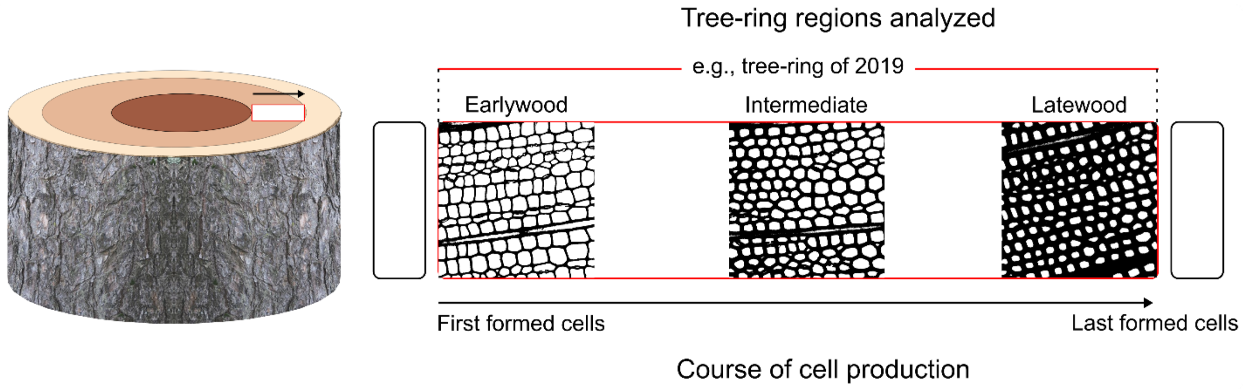

**Fig. S2** Species correlation matrices. Highlighted in yellow are the highest correlations (r-squared≥0.5) between pair of xylem traits, which were selected for the network analysis (igraph, R).

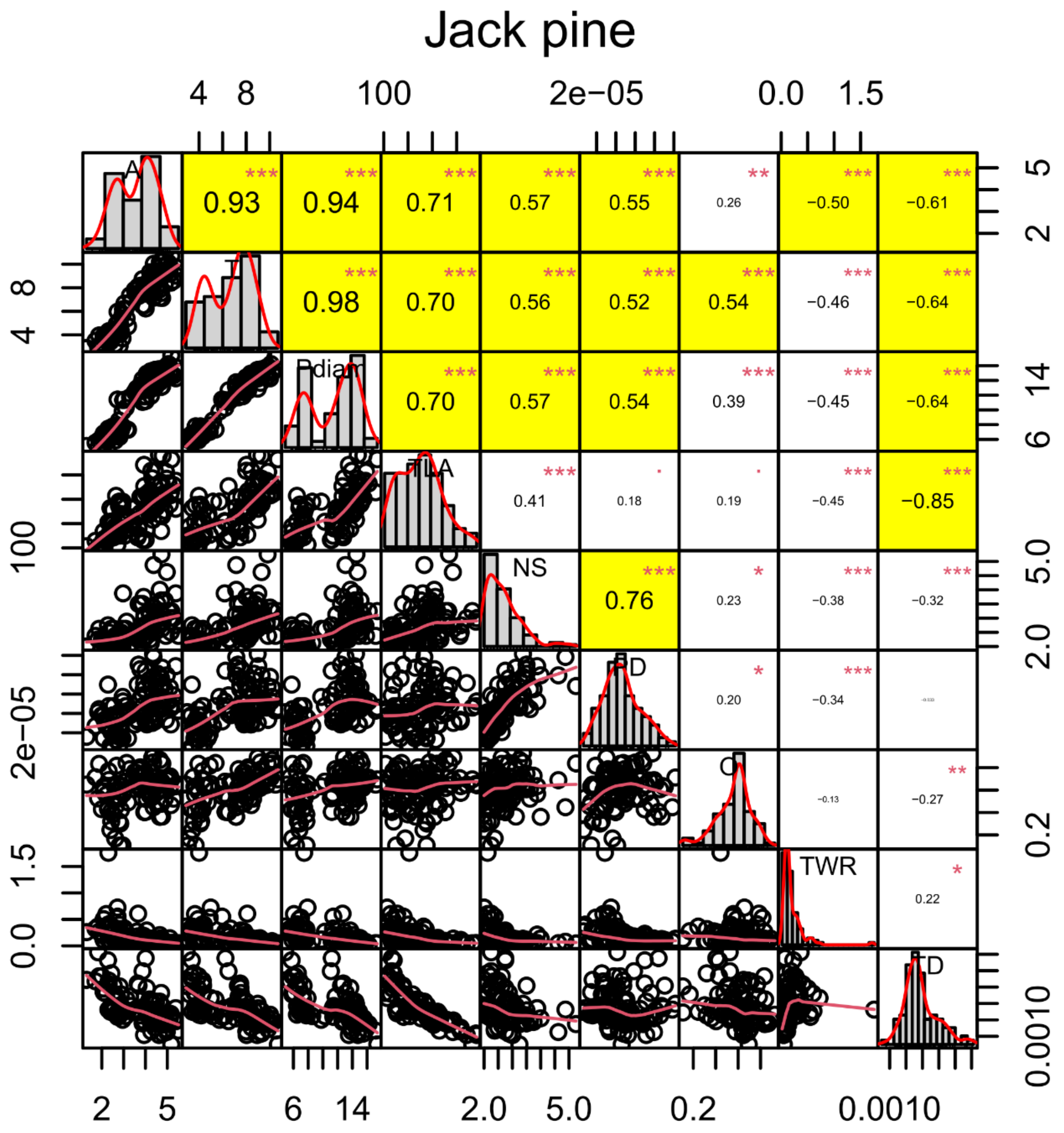

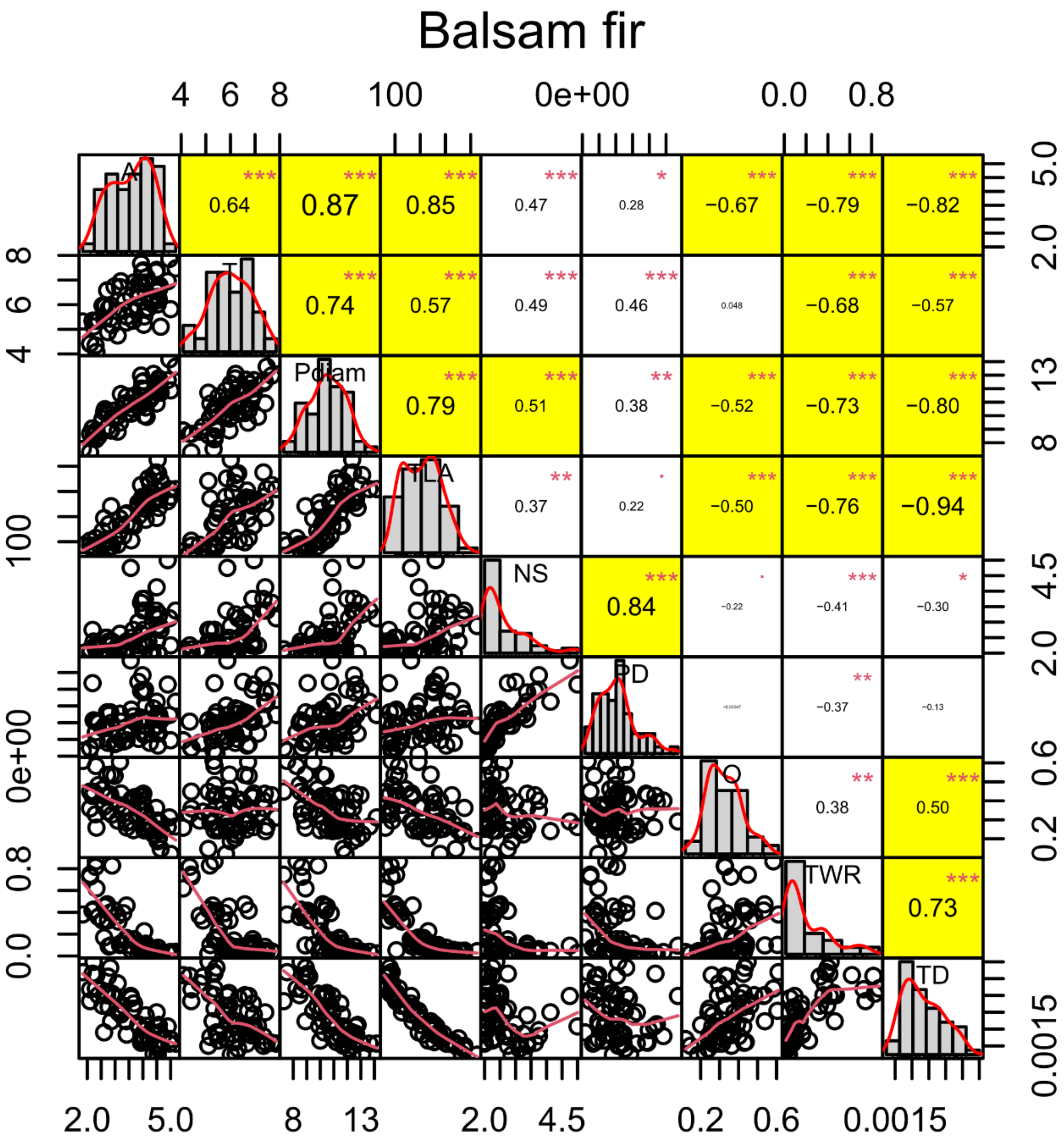

Balsam fir
A
T
Pdiam
TLA
NS
PD
TWR
TD
0.64
0.87
0.85
0.47
0.28
−0.67
−0.79
−0.82
0.74
0.57
0.49
0.46
0.048
−0.68
−0.57
0.79
0.51
0.38
−0.52
−0.73
−0.80
0.37
0.22
−0.50
−0.76
−0.94
0.84
−0.22
−0.41
−0.30
−0.37
−0.13
0.38
0.50
0.73

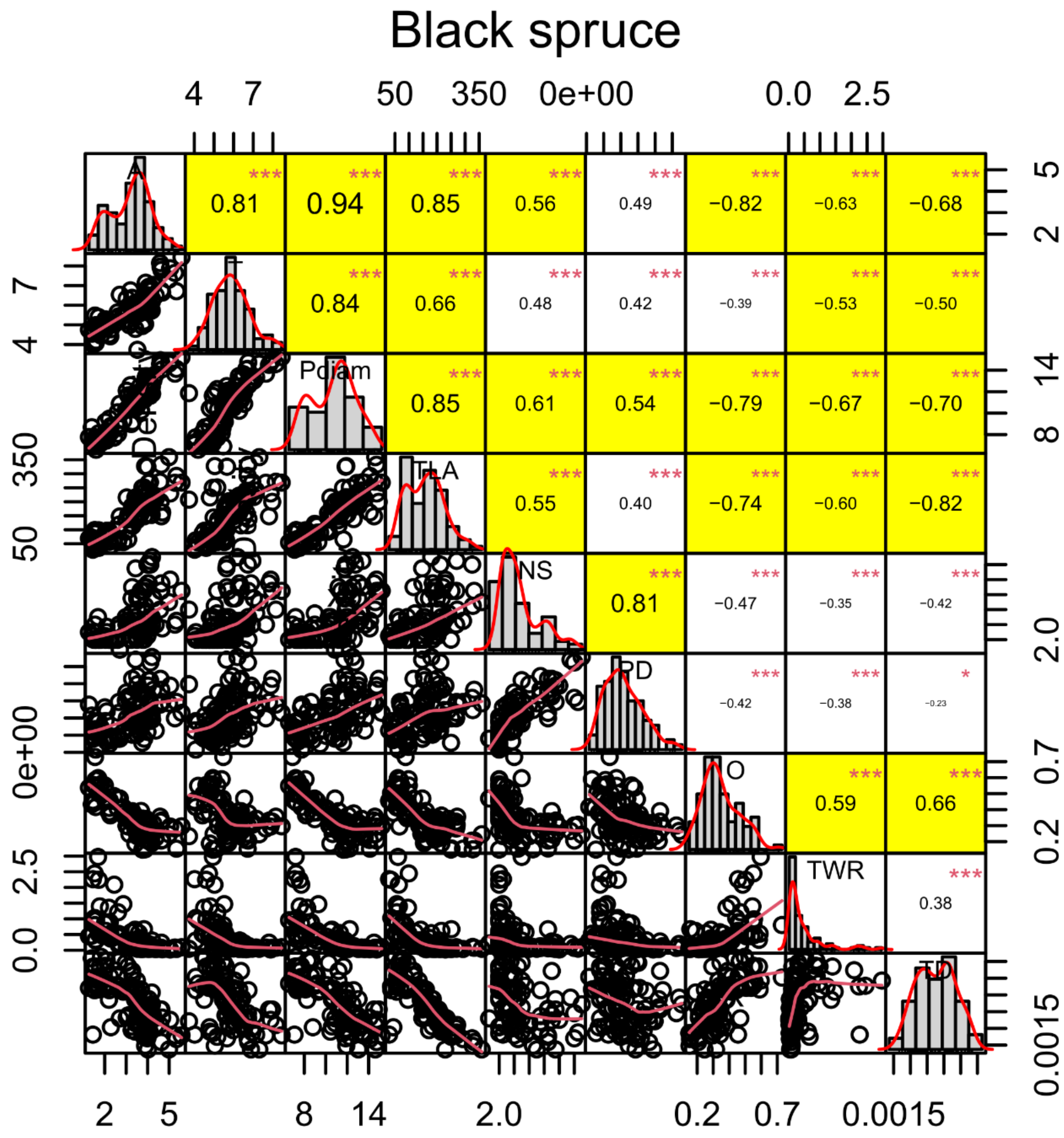

Black spruce
Pdiam
TLA
NS
PD
TWR
0.81
0.94
0.85
0.56
0.49
−0.82
−0.63
−0.68
0.84
0.66
0.48
0.42
−0.39
−0.53
−0.50
0.85
0.61
0.54
−0.79
−0.67
−0.70
0.55
0.40
−0.74
−0.60
−0.82
0.81
−0.47
−0.35
−0.42
−0.42
−0.38
−0.23
0.59
0.66
0.38
4
7
50
350
0e+00
0.0
2.5
5
2
14
8
2.0
0.7
0.2
0.0015

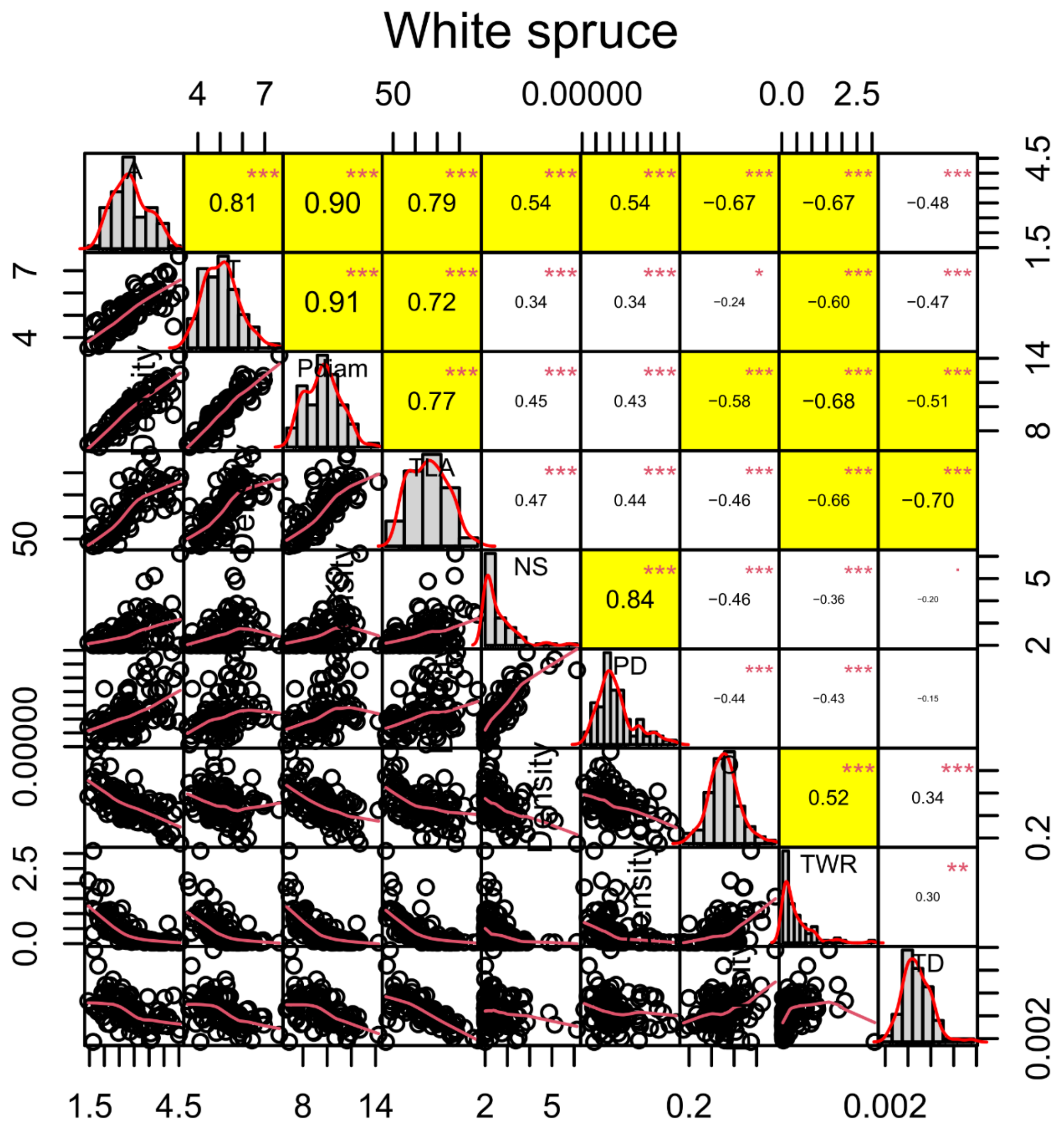
White spruce
NS
PD
TWR
TLA
0.81
0.90
0.79
0.54
0.54
−0.67
−0.67
−0.48
0.91
0.72
0.34
0.34
−0.24
−0.60
−0.47
0.77
0.45
0.43
−0.58
−0.68
−0.51
0.47
0.44
−0.46
−0.66
−0.70
0.84
−0.46
−0.36
−0.20
−0.44
−0.43
−0.15
0.52
0.34
0.30

**Fig. S3** Seasonal and annual variation in pit density (PD), tracheid density (TD), pit-tracheid net size (NS), and margo space (Msp). Letters indicate statistical differences (p-value<0.05) between species compared within the same year and growth-ring region. The test was computed via Tukey's "honest significant difference" method.

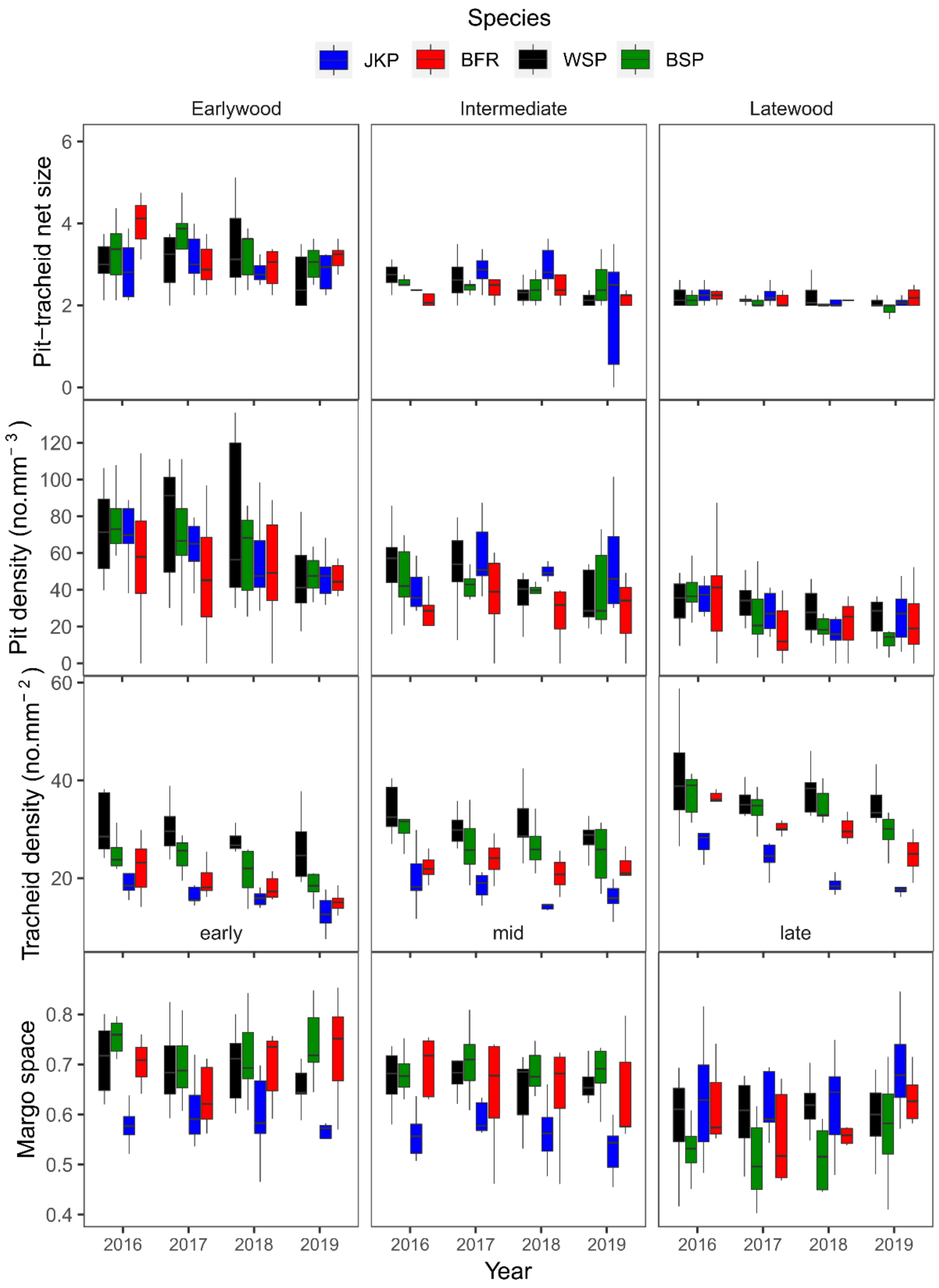

**Table S1** Regression parameters between torus diameter (Dt) and pit aperture (Da) of jack pine (JKP), black spruce (BSP), white spruce (WSP), and balsam fir (BFR), from 2016 to 2019.

| Trait interaction | Sp | 2016 | | | | 2017 | | | | 2018 | | | | 2019 | | | |
|---|---|---|---|---|---|---|---|---|---|---|---|---|---|---|---|---|---|
| | | Int | Slope | $r^2$ | p-value | Int | Slope | $r^2$ | p-value | Int | Slope | $r^2$ | p-value | Int | Slope | $r^2$ | p-value |
| Dt ~ Da | JKP | -0.08 | 1.89 | 0.93 | $2e^{-11}$ | 0.08 | 1.81 | 0.91 | $2e^{-11}$ | -1.38 | 2.20 | 0.91 | $5e^{-11}$ | -1.71 | 2.38 | 0.97 | $2.8e^{-15}$ |
| | EPN | 3.38 | 0.68 | 0.78 | $9e^{-7}$ | 3.32 | 0.74 | 0.83 | $1e^{-8}$ | 3.45 | 0.74 | 0.85 | $1.7e^{-9}$ | 3.22 | 0.82 | 0.79 | $1.1e^{-6}$ |
| | WSP | 2.70 | 0.84 | 0.81 | $1e^{-6}$ | 3.10 | 0.65 | 0.75 | $3e^{-5}$ | 2.44 | 0.95 | 0.82 | $1e^{-6}$ | 1.81 | 1.22 | 0.93 | $8.4e^{-12}$ |
| | BFR | 2.90 | 0.84 | 0.83 | $7e^{-5}$ | 4.45 | 0.52 | 0.55 | 0.017 | 3.64 | 0.68 | 0.73 | $4e^{-4}$ | 5.42 | 0.23 | 0.23 | 0.35 |
